# Outer hair cell receptor potentials reveal a local resonance in the mammalian cochlea

**DOI:** 10.1101/2023.02.24.529908

**Authors:** Andrei N Lukashkin, Ian J Russell, Oyuna Rybdylova

## Abstract

Sensory hair cells, including the sensorimotor outer hair cells, which enable the sensitive, sharply tuned responses of the mammalian cochlea, are excited by radial shear between the organ of Corti and the overlying tectorial membrane. It is not currently possible to measure directly *in vivo* mechanical responses in the narrow cleft between the tectorial membrane and organ of Corti over a wide range of stimulus frequencies and intensities. The mechanical responses can, however, be derived by measuring hair cell receptor potentials. We demonstrate that the seemingly complex frequency and intensity dependent behaviour of outer hair cell receptor potentials could be qualitatively explained by a two-degrees of freedom system with a local cochlear partition and tectorial membrane resonances strongly coupled by the outer hair cell stereocilia. A local minimum in the receptor potential below the characteristic frequency is always observed at the tectorial membrane resonance frequency which, however, might shift with stimulus intensity.

## INTRODUCTION

The mammalian cochlea is an impressively sensitive, sharp frequency analyser which works over a wide range of sound pressure levels (SPLs) exceeding six orders of magnitude (Robles and Ruggero, 2001). These features are associated with a process called the cochlear amplifier (Davis, 1983), which amplifies and sharpens cochlear responses to low-level sound stimulation but compresses them at mid to high stimulus levels. Cochlear amplification is observed only in healthy cochleae and it vanishes once cochlear function is compromised. The cellular basis of cochlear amplification is the sensory motile outer hair cells (OHCs) (Figure 1A) (Brownell et al., 1985; Liberman et al., 2002; Ashmore, 2008; Dallos, 2008; Mellado Lagarde et al., 2008). OHCs are mechanical effectors that change their length in response to changes in their transmembrane voltage (Ashmore 1987; Santos-Sacchi and Dilger, 1988). Length changes of the stiff OHCs can generate forces sufficient for minimising the damping of mechanical responses of cochlear structures surrounded by fluids (Gold, 1948; Lukashkin et al., 2007b; Dong and Olson, 2013). Three rows of OHCs are imbedded in the sensory organ of Corti (OoC) sitting on top of the extracellular basilar membrane (BM) that extends the length of the entire spiral cochlea (Figure 1A). The mechanoelectrical transducer (MET) channels of the OHCs are located close to the tips of the OHC sensory organelles, stereocilia, which form hair bundles that are imbedded in the extracellular tectorial membrane (TM) that covers the OoC, with the TM inner edge being elastically attached to the bony spiral limbus (Richardson et al., 2008). The OHC hair bundles provide a stiff mechanical link between the TM and the reticular lamina (RL) at the apical surface of the OoC. During radial shear between the TM and RL, the hair bundles are rotated about their attachment to the apical surface of the OHCs (Figure 1B), which leads to modulation of the MET current and generation of intra- and extracellular receptor potentials (RP) (Russell, 2008). RP generation results in changes in the OHC transmembrane voltage and associated OHC length changes. Hair bundles of the other type of sensory cell in the OoC, the inner hair cells (IHCs), which have rich afferent innervation and provide information to the brain, are free-standing and excited by flow of fluid entrained in the subtectorial space (STS, Figure 1A) during radial shear between the TM and RL (Sellick and Russell, 1980; Patuzzi and Yates, 1997; Nowotny and Gummer, 2006). The radial shear occurs during transversal BM vibrations (Figure 1B). The BM mechanical properties are graded and BM vibrations, which propagate as travelling waves (TWs) along the BM from the high-frequency base to low-frequency apex, peak at a frequency-specific, characteristic frequency (CF) place, where most of the TW energy is dissipated (Figure 1B) (von Békésy, 1960). Thus, the BM effectively functions as a frequency analyser separating constitutive frequency components of sounds in space and time. The TWs quickly die out at a cochlear place where the BM resonance frequency *ω*_bm_ (Figure 1B) is equal to the stimulus frequency (von Békésy, 1960). The BM is not stiff enough to support the TW beyond this point.

**Figure 1.**
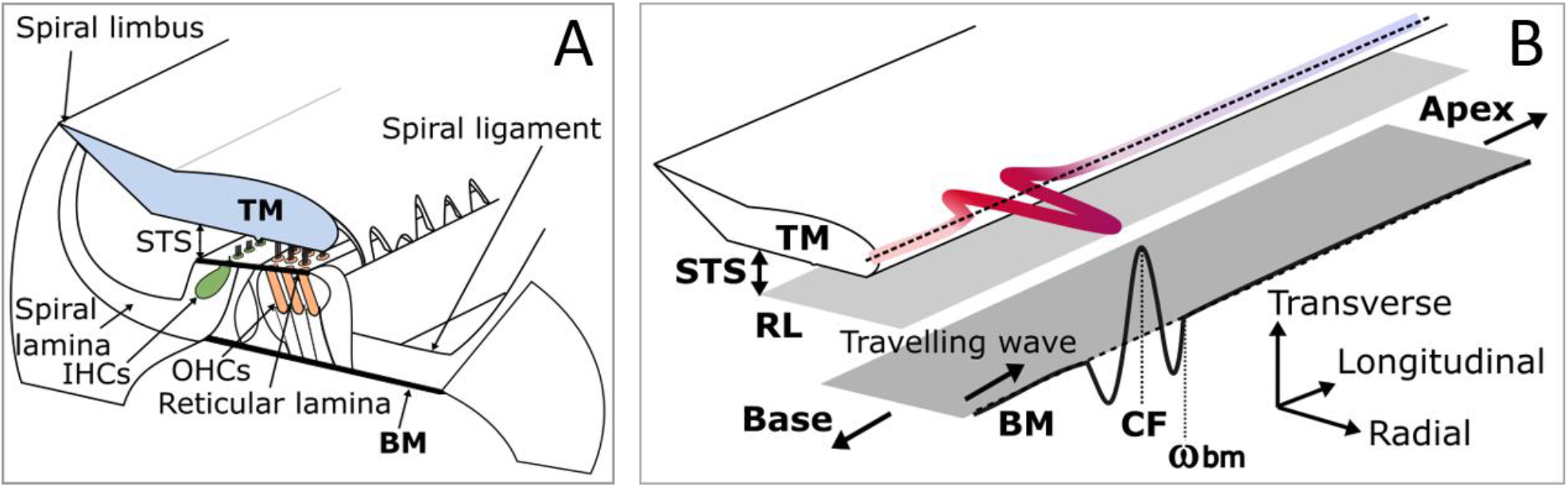
Schematics of the mammalian cochlea and the propagation of travelling wave along its length. BM, basilar membrane; TM, tectorial membrane; STS, subtectorial space; OHCs and IHCs are inner and outer hair cells, respectively; RL, reticular lamina; CF and *ω*_bm_ are the characteristic frequency place and BM resonance place for a given stimulus frequency, respectively. (A) Schematic of the cochlear cross-section showing the relationship between the BM, the sensory OoC, sitting on top of the BM, and the TM overlaying the OoC. The TM is attached to the OHC sensory stereocilia and the spiral limbus but the IHC stereocilia are free standing and deflected by flow of fluid in the STS. (B) TW propagation along the BM generates transversal movement of the BM (black wave). The TW slows down and its amplitude builds up, reaching a maximum at the CF, when the TW comes to the point where the BM resonance frequency, *ω*_bm_, is the same as frequency of the sound stimulation. The TW does not propagate beyond the *ω*_bm_ place towards the cochlear apex because the stiffness of the BM is insufficient to support the TW. The transversal BM movement is transformed into radial shearing motion between the TM and RL (red wave), which, in turn, deflects stereocilia of sensory OHCs and IHCs. Modified from (Jones et al., 2013).

The intricate structure of the cochlea (Figure 1A) appears to have evolved to enable fine-tuning of the OHCs responses and ensure optimal cochlear amplification to augment the stimulation of IHCs and, consequently, to provide adequate auditory information to the brain (Russell, 2008). Therefore, knowledge of the STS micromechanics is critical for understanding the workings of the cochlea. Current experimental methods for recording direct mechanical responses in the narrow cleft between the TM and RL do not have sufficient resolution *in vivo* to provide unambiguous data on the mechanics in this confined geometry over a wide range of stimulus frequencies and levels (Nowotny and Gummer, 2006). Fortunately, because the OHC hair bundles are embedded into the TM, and because the OHC RPs are generated due to radial shear between the TM and RL (Russell, 2008), measurement of the OHC RP can provide an insight into micromechanics of the STS. The current study demonstrates that the seemingly complex behaviour of the OHC RP recorded from a single OHC in the two-dimensional space of stimulus levels and frequencies at the cochlear base (Figure 2) (Kössl and Russell, 1992; Russell and Kössl, 1992; Levic et al., 2022) arises from a local resonance in every frequency place. A minimal mechanical arrangement, which still can explain the OHC RP behaviour, consists of a resonating BM and TM strongly coupled together via OHC stereocilia, with the TM resonance frequency located below the BM resonance frequency in every cochlear place (Allen, 1980; Zwislocki, 1980; Allen and Neely, 1992).

**Figure 2.**
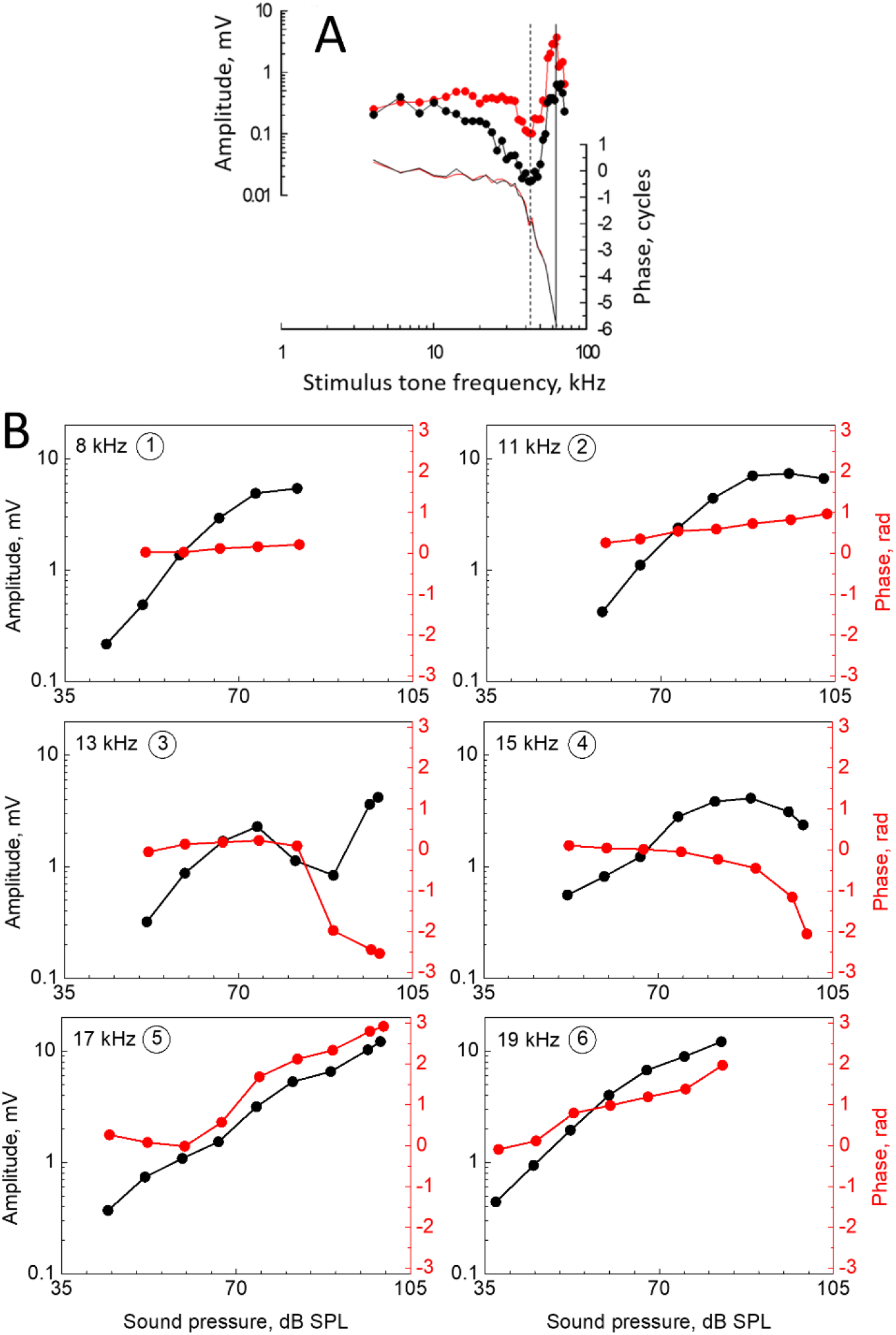
Outer hair cell receptor potentials recorded in mouse and guinea pig cochleae. (A) The 70 dB SPL isolevel frequency functions of the intracellular (black) and extracellular (red) receptor potentials recorded from the mouse cochlea. The data were compensated for recording electrode low-pass filter characteristics. Vertical solid lines indicate CF; dotted line indicate the frequency of amplitude minimum observed about half an octave below the CF. Phase was corrected for middle ear transfer characteristics (Dong et al., 2013), sound system, and recording electrode. Modified from (Levic et al., 2022). (B) Amplitude (black) and phase (red) receptor potential level functions recorded extracellularly from a guinea pig OHC (CF is 18 kHz) at the frequencies indicated within each panel. The amplitude was compensated for the single-pole, low-pass filtering of the recording electrode (corner frequency is 3.5 kHz). Numbers in circles in each panel identify frequencies with relative positions as indicated in Figure 4C. Modified from (Kössl and Russell, 1992).

## RESULTS

### Linear Passive System

A minimal micromechanical model which still qualitatively reproduces the behaviour of the OHC RP is an Allen-Zwislocki-Neely type model (Allen, 1980; Zwislocki, 1980; Allen and Neely, 1992) in which the TM is able to resonate radially due to its elastic attachments to the OHC stereocilia and the spiral limbus (Figure 1A). For this model, each cross-section of the OoC with attached TM could be represented by a schematic shown in Figure 3A (see Allen (1980) for the equivalent of this schematic and a mechanical system with the TM-RL shear).

**Figure 3.**
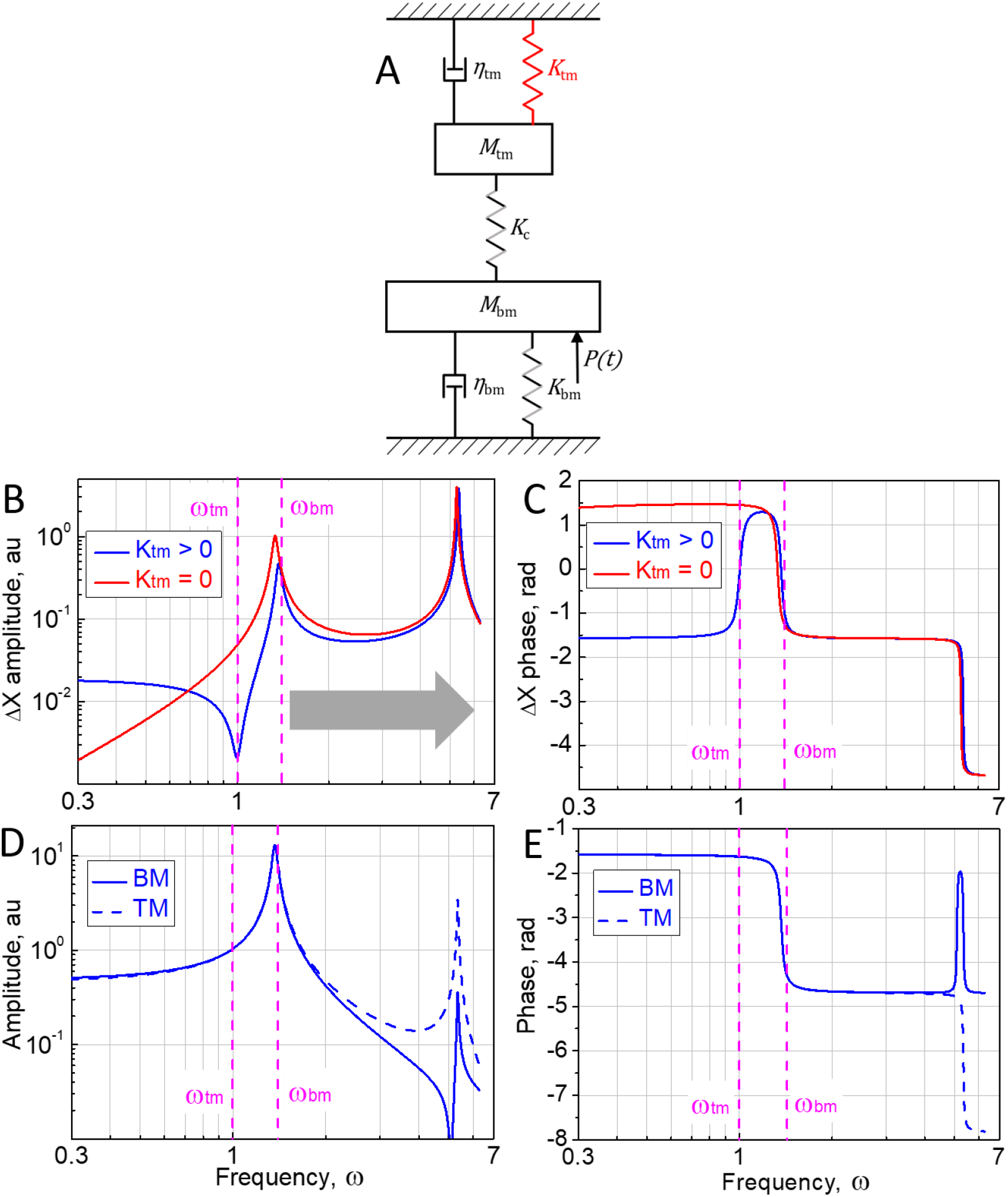
A schematic of cochlear cross-section with resonating tectorial membrane and its responses to harmonic excitation *P*(*t*). (A) A schematic showing the relationship between the mechanical elements in a cochlear cross-section. *M*_tm_ is the TM mass and *M*_bm_ denotes the entire cochlear partition mass; *K*_tm_ and *K*_bm_ are stiffnesses of the TM limbal attachment and BM stiffness respectively; *K*_c_ is elastic coupling between the TM and BM due to OHC stereocilia; *η*_tm_ and *η*_bm_ denote viscous damping of the TM and BM respectively. See the main text for more details. (B) Amplitude and (C) phase responses for the relative displacement, *ΔX*, between the BM and TM. (D) Amplitude and (E) phase responses of the BM and TM. Vertical dashed magenta lines indicate *ω*_tm_ and *ω*_bm_ as defined by equation (3). Responses for the condition *K*_tm_ = 0 in panels (D) and (E) are not shown because at the given resolution they are superimposed with responses when *K*_tm_ > 0. Horizontal grey arrow in panel (B) indicates the frequency range where *in vivo* responses are not recorded because the BM TW does not propagate beyond the *ω*_bm_ place towards the cochlear apex (Figure 1B). The following parameters were used to calculate the responses: *M*_tm_ = 1, *M*_bm_= 10, *ω*_tm_ = 1, *ω*_bm_ = 1.4, *ω*_c_ = 5, *ζ*_tm_ = *ζ*_bm_= 0.05, *P*_*a*_ = 10.

The system in Figure 3A is described by the following equations

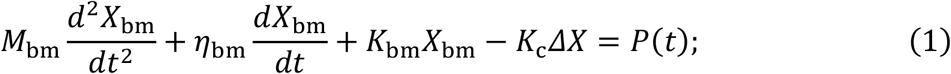

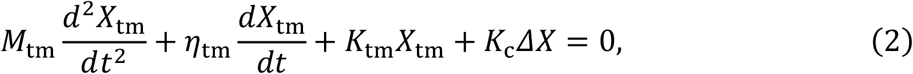

where a harmonic external force *P*(*t*) = *P*_*a*_sin(*ωt*) of frequency *ω* and amplitude *P*_*a*_ is applied to the BM, and *M*_bm_ denotes the entire cochlear partition mass. *ΔX* = *X*_tm_ − *X*_bm_ is the relative displacement between the OoC and TM which excites the OHCs and, as a first approximation for small *ΔX*, could be used to make estimates of the OHC RP behaviour.

Substituting the characteristic parameters,

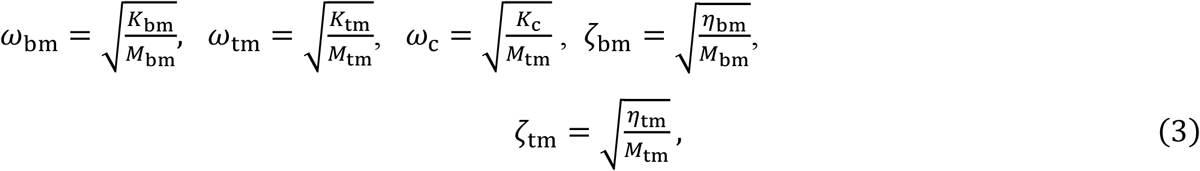

the system of equations (1, 2) may be rewritten as

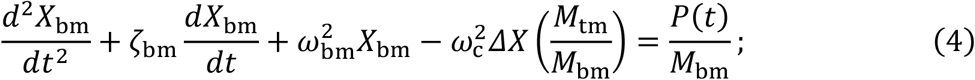

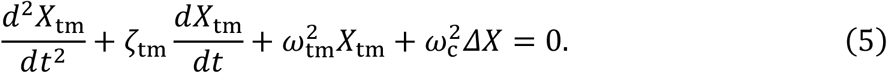

We refer to *ω*_bm_ and *ω*_tm_ as the BM and TM resonance frequency, respectively, for the rest of the paper. For the chosen relationships between the model parameters, which are in line with those measured experimentally and used in other modelling studies (e.g. see Meaud and Grosh, 2014; Nankali et al., 2022), *ΔX*, i.e. relative displacement between *M*_tm_ and *M*_bm_, demonstrates a local minimum at frequency *ω*_tm_ (Figure 3B). The minimum should result in a decrease in OHC excitation and, in turn, in a local minimum in the OHC RP observed about half an octave below the CF (Figure 2A). The minimum always occurs at the TM resonance frequency *ω*_tm_, where the TM has a minimal impedance determined only by the viscous damping and, hence, minimal load on the OHC stereocilia, and its frequency position does not depend on the properties of the driven oscillator, i.e. the BM (see question 1 and an answer to it in the Supplemental Information for detailed derivation). The *ΔX* minimum becomes more pronounced with decreasing TM damping so that *ΔX* → 0 at frequency *ω*_tm_ when *ζ*_tm_ → 0.

For the chosen model parameters, the minimum/antiresonance is not observed in the BM and TM responses for frequencies below *ω*_bm_ (Figure 3D). Δ*X* is stiffness dominated below the minimum (Figure 3C). Δ*X* becomes mass dominated at the amplitude minimum and a corresponding phase transition of *π* is observed (also see Equation S24 in the Supplemental Information). The corresponding phase transition is also observed near the minimum of the experimentally measured OHC RP (Figure 2A). Δ*X* becomes stiffness dominated close to the first normal mode of vibrations near *ω*_bm_ where local maximums of the BM and TM displacements (Figure 3D) and Δ*X* (Figure 3B) are observed, and where the Δ*X* phase angle returns to − ^*π*^/_2_ (Figure 3C, Equation S24 in the Supplemental Information). It should be noted that the system of equations (4, 5) does not include the TW observed in the cochlea. Therefore, a large phase roll-off due to TW propagation (Figure 2A) is not observed in the model responses. The phase demonstrates only a transition up to 180 degrees (Figure 3C) for the mass-dominated responses between *ω*_tm_ and *ω*_bm_, which is similar to that seen in figure 3 of Allen (1980). Also, because the model does not include TW, a sharp decline in the amplitude of the OHC RP above the CF (Figure 2A) is not observed in the model presented in Figure 3A. In the real cochlea, the TWs quickly die out at a cochlear place where the BM resonance frequency *ω*_bm_ is equal to the stimulus frequency (Figure 1B) and responses for frequencies above *ω*_bm_ cannot be recorded (von Békésy, 1960). This frequency region is indicated by horizontal grey arrows in Figures 3B, 4B. Hence, the second normal mode, which is shifted to frequencies well above *ω*_bm_ due to strong elastic coupling *K*_c_ between *M*_tm_ and *M*_bm_, is not observed *in vivo* (see question 2 and an answer to it in the Supplemental Information for detailed derivation).

The role of the TM limbal attachment *K*_tm_ for OHC excitation was investigated experimentally by Lukashkin et al. (2012) and modelled by Meaud and Grosh (2014). Similar sensitivity and sharpness of BM tuning were found in wild-type mice and mutant mice with the TM detached from the spiral limbus. It was suggested that the elasticity of the TM attachment to the spiral limbus is not a crucial factor for exciting the OHCs near their CF, and that the OHCs must be excited by the inertial load provided by the TM mass at CF to effectively boost the mechanical responses of the cochlea. Indeed, while the Δ*X* minimum is not observed in model responses when *K*_tm_ = 0 (Figure 3B, see question 3 and an answer to it in the Supplemental Information for detailed derivation), Δ*X* response is mass dominated and the phase angle is similar in both cases, *K*_tm_ > 0 and *K*_tm_ = 0, between *ω*_tm_ and *ω*_bm_ (Figure 3C). In the real cochlea, this frequency range corresponds to stimulus frequencies below the CF where the non-linear cochlear amplification gradually builds up (Nilsen and Russell, 1999; Robles and Ruggero, 2001; Zheng et al., 2007; Dong and Olson, 2013; Lee et al., 2016) and, thus, the Δ*X* phase angle and OHC excitation timing are optimal for cochlear amplification to occur. Therefore, similar Δ*X* excitation phase/timing for conditions *K*_tm_ = 0 and *K*_tm_ > 0 (Figure 3C), with *K*_tm_ = 0 simulating mutants with the TM detached from the spiral limbus, supports the conclusion that the OHCs must be excited by the inertial load provided by the TM mass at CF to effectively boost the mechanical responses of the cochlea (Gummer et al., 1996; Lukashkin et al., 2010; Lukashkin et al., 2012; Meaud and Grosh, 2014; Nankali et al., 2022).

### Active Nonlinear System

The cochlear amplifier is introduced as a nonlinear damping (*η*_n_ in Figure 4A) which includes a level-independent positive damping and a level-dependent negative damping component (Gold, 1948; Elliott et al., 2015) demonstrated experimentally (Lukashkin et al., 2007b) so that

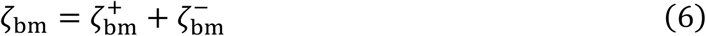

and

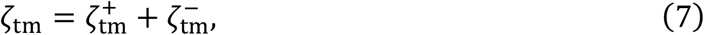

where *ζ*^+^ and *ζ*^−^ are corresponding positive and negative components for the BM and TM damping. The cochlear amplifier emerges from the OHC length changes which are controlled by changes in the voltage across the OHC basolateral membrane (Ashmore 1987; Santos-Sacchi and Dilger, 1988). The transmembrane voltage changes, in turn, are generated by the MET current which is modulated when the OHC hair bundles are rotated about their attachment to the OHC apical cuticular plate due to the relative displacement between the OoC and TM (Russell, 2008). In this case, the OHC MET current is a function of *ΔX*. As a first approximation this function could be describes by a sigmoidal nonlinearity/Boltzmann function

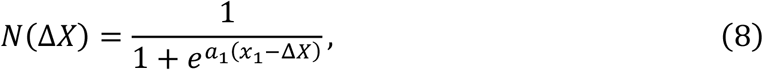

where *N*(Δ*X*) is the MET nonlinearity, *a*_1_ sets the slope of the function, and *x*_1_ is the position of maximum slope. Therefore, the negative damping for both the BM and TM could be defined as

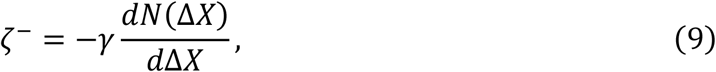

where *γ* is the transfer ratio relating the change in the OHC receptor potential to resultant negative damping developed by the OHCs. To find numerical solutions of equations (4-9) in the time domain, *γ* was taken to be *γ* = *4*(*ζ*^+^ − 0.0001) for both the BM and TM to ensure that the total damping is always positive.

**Figure 4.**
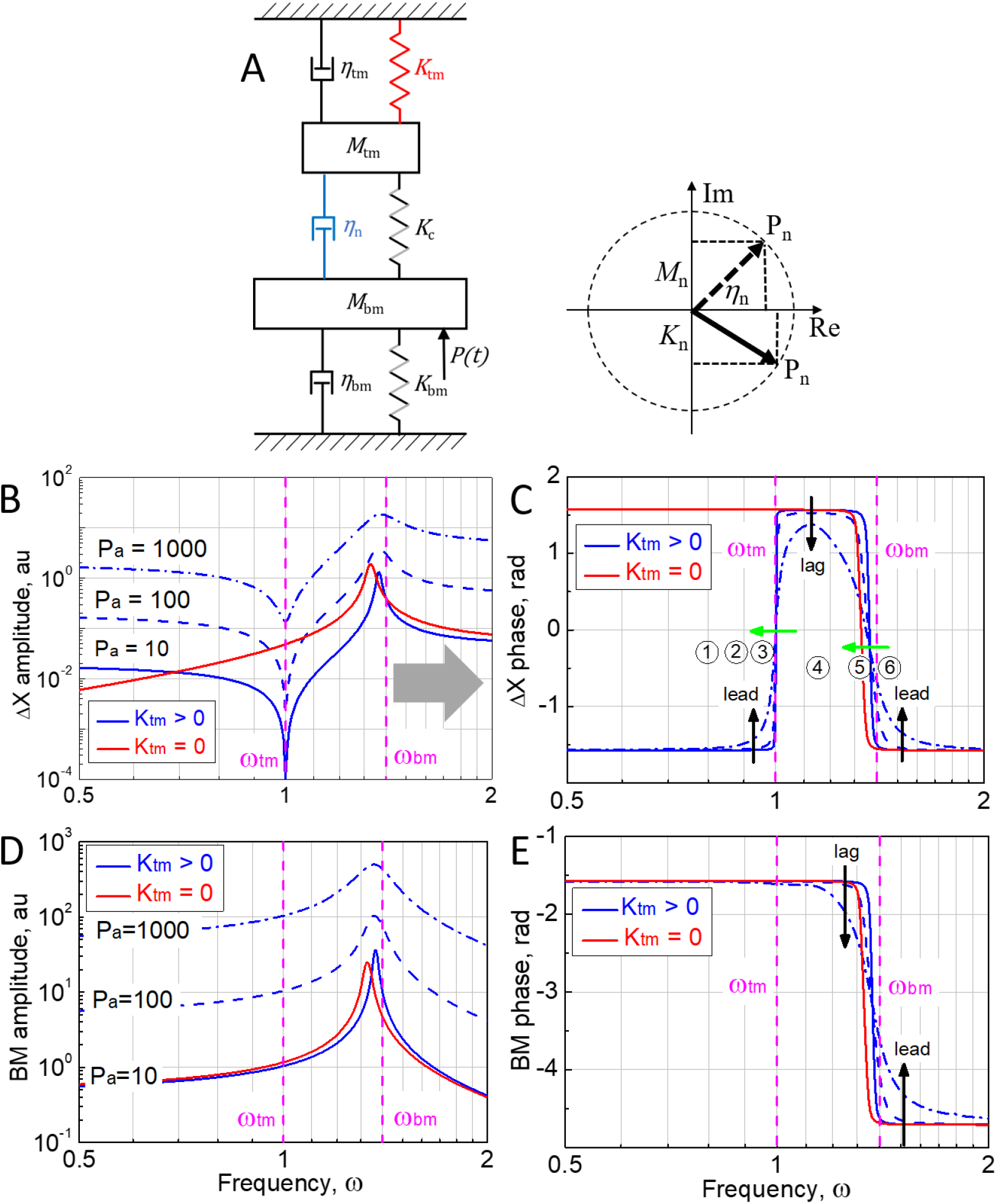
Response of the active model with nonlinear damping to harmonic excitation. (A) A schematic showing the relationship between the mechanical elements in a cochlear cross-section. *M*_tm_ is the TM mass and *M*_bm_ denotes the entire cochlear partition mass; *K*_tm_ and *K*_bm_ are stiffnesses of the TM limbal attachment and BM stiffness respectively; *K*_c_ is elastic coupling between the TM and BM due to the OHC stereocilia; *η*_tm_ and *η*_bm_ denote viscous damping of the TM and BM respectively; *η*_n_ is a nonlinear damping due to action of the OHCs. Only *P*_*a*_ = 10 response for *K*_tm_ = 0 is shown. Insert in (A) illustrates a situation when the nonlinear, level-dependent OHC force *P*_n_ is out of phase with the damping force *η*_n_. In this case a nonlinear, level-dependent stiffness *K*_n_ (if *P*_n_ lags *η*_n_) or mass *M*_n_ (if *P*_n_ leads *η*_n_) associated with the OHC action is observed. See the main text for more details. (B) Amplitude and (C) phase responses of the relative displacement, *ΔX*, between the BM and TM for different amplitudes *P*_*a*_ of the harmonic force *P*(*t*). *P*_*a*_ is indicated for each curve. Responses for the condition *K*_tm_ = 0 are shown only for *P*_*a*_ = 1. Vertical black arrows in (C) show changes in the *ΔX* phase with increasing *P*_*a*_, i.e. the stimulus level. Numbers in circles in (C) identify presumed frequency positions of the corresponding experimental responses (Figure 2B) relative to *ω*_tm_ and *ω*_bm_. Green horizontal arrows in (C) shows a presumed shift of *ω*_tm_ and *ω*_bm_ to lower frequencies due to changes in *K*_n_. See inset in (A). (D) Amplitude and (E) phase angle of BM responses for different amplitudes *P*_*a*_ of the harmonic force *P*(*t*). *P*_*a*_ is indicated for each curve. Responses for the condition *K*_tm_ = 0 are shown only for *P*_*a*_ = 1. Vertical black arrows in (E) show changes in the BM phase responses with increasing *P*_*a*_, i.e. the stimulus level. Vertical dashed magenta lines indicate *ω*_tm_ and *ω*_bm_ as defined by equation (3). A horizontal grey arrow in (B) indicates the frequency range where *in vivo* responses are not recorded because the BM TW does not propagate beyond the *ω*_bm_ place towards the cochlear apex (Figure 1B). The following parameters were used to calculate the responses: 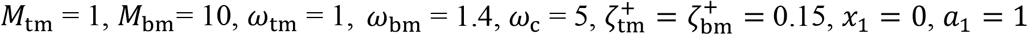.

Responses of the active model to harmonic excitation *P*(*t*) = *P*_*a*_sin(*ωt*) for different amplitude *P*_*a*_ are shown in Figure 4. Only responses below and around *ω*_bm_ are shown because the BM TW does not propagate beyond the *ω*_bm_ place towards the cochlear apex (horizontal grey arrow in Figure 4B). Therefore, the second normal mode (see question 2 and an answer to it in the Supplemental Information for detailed derivation), which is shifted to frequencies well above *ω*_bm_ due to strong elastic coupling *K*_c_ between *M*_tm_ and *M*_bm_, is not shown in Figure 4 (compare Figure 3B and 4B). An active model, which includes only local BM/TM resonances, provides an impressively good qualitative description of the experimental data for the OHC receptor potentials (Figure 2) despite the absence of global phenomena like the BM TW or elastic/hydromechanical coupling along the cochlea. The RP minimum seen about half an octave below the CF in isolevel RP responses recorded for a stimulus level of 70 dB SPL to ensure the recording of responses over a wide frequency range (Figure 2A) is less sharp than the model minima recorded for smaller *P*_a_ (Figure 4B). However, the model minimum becomes less sharp for *P*_a_ = 100 when the nonlinear model amplification becomes saturated (see question 1 and an answer to it in the Supplemental Information for the assessment of the depth of the local minimum at *ω*_tm_ for varying TM damping).

The active local resonance model provides an explanation of the seemingly complex level-dependent OHC RP amplitude and phase behaviour observed in experiments (Figure 2B, Kössl and Russell, 1992) assuming that *ω*_tm_ was situated around 13 kHz, i.e. about half an octave below the CF of 18 kHz. Indeed, in this case, the phase angle does not depend on the stimulus level when the stimulus frequency is 8 kHz and well below *ω*_tm_ (panel 1 in Figure 2B and frequency point 1 in Figure 4C). There is a small phase lead with level (leftmost black vertical arrow in Figure 4C) for the stimulus frequency of 11 kHz which is situated closer to but still below *ω*_tm_ (panel 2 in Figure 2B and frequency point 2 in Figure 4C) but the phase lead is larger (rightmost black vertical arrow in Figure 4C) for the stimulus frequency of 19 kHz above the CF/*ω*_bm_ (panel 6 in Figure 2B and frequency point 6 in Figure 4C). The phase behaviour reverses and the phase lags with level (middle vertical black arrow in Figure 4C) for the stimulus frequency of 15 kHz situated between the *ω*_tm_ and CF/*ω*_bm_ (panel 4 in Figure 2B and frequency point 4 in Figure 4C). A reversal of phase behaviour is observed with increasing the stimulus level for 13 kHz (small lead to lag) situated just below assumed *ω*_tm_, and 17 kHz (small lag to lead) situated just below the CF/*ω*_bm_ in panels 3 and 5 in Figure 2B, respectively. At the same time, this reversal of the phase behaviour for both stimulus frequencies are associated with phase transitions which are close to 180°. The near-180° phase transition at 13 kHz is steeper than the transition observed for 17 kHz which is expected because of a steeper phase slope near *ω*_tm_ (Figure 4C). An amplitude notch in the OHC RP level function is, however, observed only for 13 kHz stimulus (panel 3 in Figure 2B) and it is absent at 17 kHz. In terms of the active model (Figure 4A), the observed nonmonotonic amplitude and phase behaviour is explained by presumed shifts of *ω*_tm_ and *ω*_bm_ to lower frequencies with increasing the stimulus level (green horizontal arrows in Figure 4C). Indeed, the low-frequency shift of maximum responses near the CF is well-documented (e.g. Robles and Ruggero, 2001) and a low-frequency shift of the TM resonance was suggested from observation of different indices of cochlear responses associated with *ω*_tm_ (Lukashkin et al., 2007a). Therefore, if 13 kHz (panel 3 in Figure 2B) is situated just below *ω*_tm_ (frequency point 3 in Figure 4C) for low stimulus levels but it appears above *ω*_tm_ for high stimulus levels then the *ΔX* amplitude (i.e. the OHC RP amplitude) would fall into the amplitude minimum at *ω*_tm_ and eventually recover from it with increasing stimulus level. In this case not only a reversal of the phase behaviour and a steep phase transition but also an amplitude notch in the OHC RP level functions should be observed. The amplitude notch in the OHC RP level functions should not be observed during reversal of the phase behaviour and corresponding phase transition for responses to the 17 kHz stimulus (panel 5 in Figure 2B) if this stimulus frequency is situated just below the CF/*ω*_bm_ (frequency point 5 in Figure 4C) at low stimulus levels but appears above CF/*ω*_bm_ for high stimulus levels due to a low-frequency shift of *ω*_bm_ because there is no a local amplitude minimum associated with the *ΔX* frequency responses near *ω*_bm_ (Figure 4B).

A local resonance active cochlear model which includes only nonlinear negative damping (Figure 4A) does not reproduce the suggested low-frequency shift of *ω*_tm_ and *ω*_bm_ at high stimulus levels (green arrows in Figure 4C). The low-frequency shift of the TM and BM resonances would occur quite naturally if the nonlinear, level-dependent OHC force *P*_n_ is out of phase with the damping force *η*_n_ (Figure 4A, insert). In this case nonlinear, level-dependent stiffness *K*_n_ or inertial *M*_n_ projections associated with the OHC action that correspond to the imaginary parts of the impedance are observed for mechanical components of the system (e.g. see Kolston et al., 1990). Changes in the projections *K*_n_ or *M*_n_ due to changes in the phase angle of *P*_n_ or variation of its amplitude with increasing stimulus level, would lead to changes in the imaginary part of the components’ impedances, thus changing frequencies *ω*_tm_ and *ω*_bm_. Changes in the effective masses or stiffnesses of the system components, and, hence, shifts of *ω*_tm_ and *ω*_bm_, might also be explained by a spread of excitation along the cochlea due to stiffening of the TM or/and entraining larger masses of cochlear fluids with increasing stimulus levels (see Discussion for more details). Also, the shift may be a product of OHC efferent activation (Guinan, 2018) but it should still be associated with changes in the imaginary part of the impedances, i.e. the effective stiffness/mass changes, even in this case.

It is worth noting that the minimal local resonance active cochlear model with negative damping (Figure 4A) also qualitatively reproduces experimental level dependent behaviour of the BM phase. It has been known for a long time that the phase angle of BM responses lags/leads with levels for frequencies below/above the CF/*ω*_bm_, respectively (Robles and Ruggero, 2001). Exactly this type of phase behaviour is observed for the local resonance active cochlear model with negative damping (black vertical arrows in Figure 4E).

## DISCUSSION

The objective of this study is to find a minimal mechanical system which still can qualitatively explain the behaviour of the OHC RP *in vivo*. It is demonstrated that a local resonance of a strongly coupled TM and BM as suggested by (Allen, 1980; Zwislocki, 1980; Allen and Neely, 1992) and experimentally recorded by (Gummer et al., 1996; Lee et al., 2016) is sufficient to explain the phenomenology of the seemingly complex changes in the OHC RP amplitude and phase recorded close to OHCs *in vivo* for wide range of frequencies and levels of acoustic stimulation (Kössl and Russell, 1992; Russell and Kössl, 1992; Levic et al., 2022). Moreover, the model reveals that the nonmonotonic amplitude behaviour (i.e. local minima/maxima) of experimentally recorded cochlear responses generated due to radial shear between the TM and OoC at frequencies below the CF, is observed at the TM resonance frequency and the frequency position of these minima/maxima does not depend either on the properties of the pressure driven part of the system (i.e. the BM with the OoC sitting on its top) or the degree of coupling between the TM and OoC. The model confirms that the shear between the TM and OoC is mass dominated in the frequency region associated with nonlinear cochlear amplification and a corresponding phase transition is observed for this frequency region between *ω*_tm_ and *ω*_bm_ (Figure 3C, 4C), where the timing of OHC stimulation is optimal for cochlear amplification. An obvious conclusion is that the timing is suboptimal for cochlear amplification and sharpened frequency tuning of mechanical responses of the cochlear partition outside of the range of between *ω*_tm_ and *ω*_bm_, despite the finding that OHCs are stimulated over a wider frequency span than [*ω*_tm_, *ω*_bm_] as judged by the reticular lamina active responses (Ren et al., 2016; He et al., 2022).

There is no need for global phenomena, e.g. TW or elastic/hydromechanical coupling along the cochlea, to explain the experimental data on the OHC RP (Figure 2B). In fact, addition of global phenomena to the local resonance model can smear sharp antiresonance/resonance in the system as discussed below. This, however, does not destroy the effect of local resonance. Its signature could still be seen in various types of cochlear responses (Lukashkin et al., 2010). Addition of the TW to the local resonance model, which includes independent TM resonance at frequencies below the BM resonance, provides a good fit to the neural data (Allen, 1980; Allen and Neely, 1992; Allen and Fahey, 1993). A notch of insensitivity seen in the neural data at a frequency about half an octave below the CF (Liberman, 1978; Allen, 1980; Liberman and Dodds, 1984; Taberner and Liberman, 2005; Temchin et al., 2008) resembles a similar notch in the OHC RP (Figure 2A) and in neural suppression tuning curves (Lukashkin et al., 2007a). The notch disappears in mutants with the TM detached from the spiral limbus (Lukashkin et al., 2012), confirming its origin (compare with the red curve for *K*_tm_ = 0 in Figure 3B). It was also suggested that the amplitude and phase dependence of the distortion product otoacoustic emission (DPOAE) on the ratio of the two primary stimulus tones (f1 and f2, f2 > f1) used to evoke the DPOAEs, reflected band-pass filtering of the DPOAEs by the mechanical filter associated with the local TM resonance. Amplitude maxima for DPOAEs of different order (i.e. 2f1-f2, 3f1-2f2, 4f1-3f2) are observed at the same frequency which is independent of the f2/f1 ratio (Brown et al., 1992; Allen and Fahey, 1993) and the phase of DPOAE of different order changes from lag to lead at the same frequency when the levels of primaries are increased (Lukashkin and Russell, 2003; Lukashkin et al., 2007a).

The local minimum in sensitivity of the OHC RP at frequencies about half an octave below the CF (Figure 2A) and corresponding amplitude notches associated with steep phase transitions (panel 3 in Figure 2B) are more sharply tuned in intracellular OHC RP recordings or extracellular recordings in the closest vicinity of OHCs (Kössl and Russell, 1992; Russell and Kössl, 1992; Levic et al., 2022) than when measured from the OoC fluid space (Fridberger et al., 2004) and close to the BM as a cochlear microphonic (Dong and Olson, 2013), when it becomes increasingly broader and less distinct with increasing stimulation levels. We attribute this difference to two effects, namely, to the level-dependent increase in the damping as illustrated in Figure 4B and to the level-dependent increase in the numbers of generators (OHCs) contributing to the extracellular signal (Patuzzi et al., 1989) that smears the phase data, rather than to the single effect of fluid damping, as has been recently suggested and modelled based on cochlear microphonic measurements (Nankali et al., 2020). The same level-dependent summation of electrical signals from a gradually increasing number of generators leading to a partial phase cancellation might explain lack of an obvious low-frequency shift of the minimum in the gross OHC electrical responses (Fridberger et al., 2004; Dong and Olson, 2013), the shift was suggested to explain the level dependent behaviour of different indices of cochlear responses associated with the TM resonance (Lukashkin et al., 2007a) and the OHC RP data (green arrows in Figure 4C).

It was suggested that a low-frequency shift of *ω*_tm_ and *ω*_bm_ (green arrows in Figure 4C), and corresponding shift of the low-frequency minimum of OHC RP and its local maximum near the CF with increasing the sound intensity might occur due to the desynchronization of the nonlinear, level-dependent OHC force *P*_n_ contributing to the imaginary parts of the mechanical impedances of the components (insert in Figure 4A). There are two additional mechanisms which might contribute to the low-frequency shift. Vibration of an individual element within the cochlea generates a near field pressure, which increases the element’s effective mass (Ni and Elliott, 2015). Increase in the TW wavelength with sound intensity might increase the fluid-loaded mass on the individual elements (Steele and Taber, 1979; Elliott et al., 2022; Nankali et al., 2022), thus, lowering their resonance frequencies (Equation 3). Also, the TM material properties are frequency and, thus, velocity dependent and the TM shear storage modulus increases with stimulation frequency/velocity, especially at the cochlear base (Jones et al., 2013). Increase in the TM velocity response with stimulus level and resultant TM stiffening might lead to larger regions of the TM and OoC being involved in local OHC excitation due to increased elastic coupling along the TM (Dewey et al., 2018). This would manifest in larger *M*_tm_ and *M*_bm_, and corresponding decrease in *ω*_tm_ and *ω*_bm_.

It should be noted that the OHC RP data which are analysed in this study were obtained from the high-frequency cochlear base. Both the mechanical (Recio-Spinoso and Oghalai, 2017; Burwood et al., 2022) and neural (Liberman, 1978; Allen, 1980; Liberman and Dodds, 1984; Taberner and Liberman, 2005; Temchin et al., 2008) responses at the extreme low-frequency cochlear apex have much broader tuning and lack a low-frequency shoulder, and neural responses have no local sensitivity minimum below the CF. In fact, a large stretch of the cochlear partition at the extreme apex moves in phase, making phase-locking of neural responses (Rose et al., 1967; Kim and Molnar, 1979; Johnson, 1980; Palmer and Russell, 1986) a preferable mechanism of frequency coding in this cochlear region. The difference in responses between the base and apex possibly reflects the relative difference in mechanical properties of cochlear structures. The dimensions of the TM vary along the length of the cochlea; its radial width and cross-sectional area and, hence, linear mass density increase from the basal to the apical end of the cochlea (Richardson et al., 2008). The lengths of the OHC hair bundles increase from the cochlear base to apex (Wright, 1984; Yarin et al., 2014), which results in a decrease of their rotational stiffness (Tobin et al., 2019; Miller et al., 2021) and, thus, reduction in elastic coupling between the TM and OoC.

### Limitations of the study

The objective of this work is to find a minimal model which still can qualitatively explain the complex behaviour of the OHC RP for different levels and frequencies of stimulation below and around the CF at the high frequency cochlear base. For this purpose, the model does not include global phenomena, e.g. TW along the BM and elastic coupling along the cochlear partition, and it cannot be used to fit experimental data. Only the qualitative behaviour of the OHC RP amplitude and phase responses is considered. The model assumes uniform negative damping/cochlear amplification over the entire frequency range, which, however, does not affect the conclusions because the conclusions are based on the OHC RP behaviour around the frequency range of [*ω*_tm_, *ω*_bm_] where nonlinear cochlear amplification is observed. Also, the model explains OHC RPs recorded at the high-frequency cochlear region and responses at the extreme low-frequency cochlear apex might not be well explained.

## METHODS

MATLAB (The MathWorks. Inc. 2022a) was used to find solutions for the linear (Figure 3) and nonlinear (Figure 4) models in the time domain for a harmonic stimulation *P*(*t*) = *P*_a_sin(*ωt*) and the fast Fourier transform was applied to the solutions to extract the component at frequency *ω*.

## ACKNOWLEDGMENTS

This work was funded by United Kingdom Medical Research Council Grant MR/W028956/1. OR is supported by the UKRI Future Leaders Fellowship (Grant MR/T043326/1).

## AUTHOR CONTRIBUTIONS

IR and ANL conceived and designed the study. OR completed the analytical analysis of the mechanical systems. ANL performed computational simulations. All authors wrote the manuscript.

## Supplemental information

We would like to analyse the system of equations (4, 5) to answer the following questions:

1. What is the condition for a minimum of *ΔX* at *ω*_tm_ to occur?
2. What is contribution of the normal modes into the response?
3. Does the *ΔX* minimum occur when *K*_tm_ is absent, i.e. when there is no limbal attachment of the TM to the spiral limbus?

To answer the question 1, and using the complex-exponential method, the equations (4, 5) can be rewritten

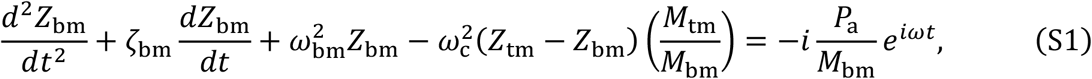

and

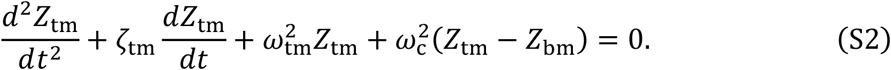

Let us assume the steady state solution in the following form:

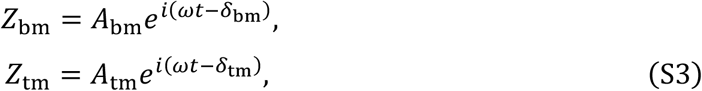

with *X*_bm_ = *Re*(*Z*_bm_) and *X*_tm_ = *Re*(*Z*_tm_).

Substituting expressions (S3) for *Z*_bm_ and *Z*_tm_ into equations (S1, S2) yields

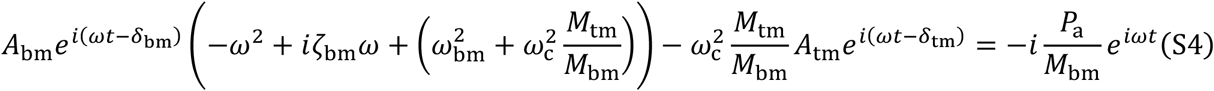

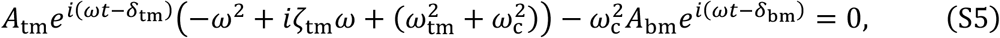

and after re-arranging

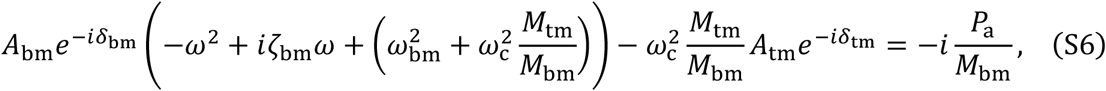

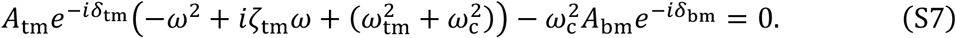

From equation (S7), we have

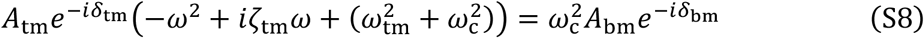

or

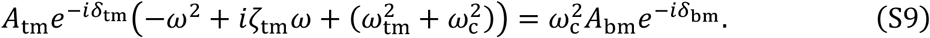

Denote 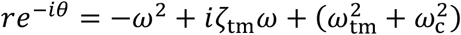, where 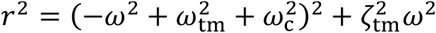 and

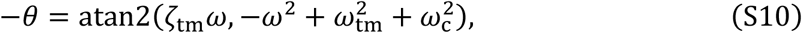

where atan2 is a four-quadrant inverse tangent, then

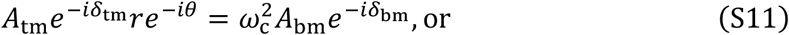

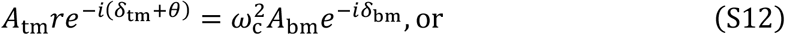

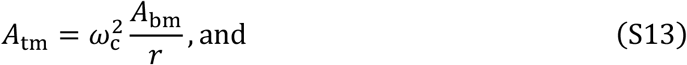

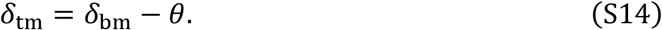

Local minimum of *ΔX* is observed when the amplitudes and phases of *Z*_bm_ and *Z*_tm_ are close. Let us compare pairs *A*_tm_ and *A*_bm_, *δ*_tm_ and *δ*_bm_ in (S13, S14). In case of light damping, *ζ*_bm_, *ζ*_tm_ ≪ *ω*_bm_, *ω*_tm_, *ω*_c_. Therefore, from the definitions (S10) of *r*, when *ω* = *ω*_tm_, *r* is close to 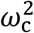 (the difference 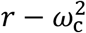 is proportional to *ζ*_tm_ ≪ 1) and from (S13) *A*_tm_ ≅ *A*_bm_. Also, since *ζ*_tm_ ≪ 1, from the definition (S10) of *θ*, when *ω* = *ω*_tm_, −*θ* is proportional to 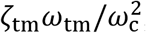, which is a small value as well, and *δ*_tm_ ≅ *δ*_bm_. To summarise, when *ω* = *ω*_tm_, amplitudes *A*_tm_ and *A*_bm_, and phases *δ*_tm_ and *δ*_bm_ of *X*_tm_ and *X*_bm_ are close, and their difference *ΔX* → 0 when *ζ*_tm_ → 0.

Note that from the definition of *θ* (S10), it follows that at 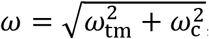, *θ* changes its value from 0 to −*π*. In other words, and if neglecting small terms, this can be rewritten as

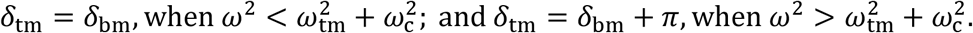

The modelling results support this outcome, see Figure 3E, where the phases of TM and BM responses coincide until 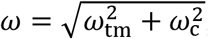, after which there is a jump in the value between the phases by *π*.

After substituting notations introduced in (S13) and (S14), to (S6), we have

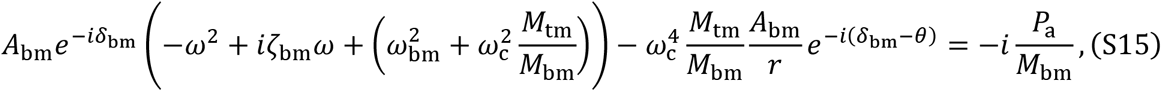

As in the analysis above, let us introduce new variables as follows

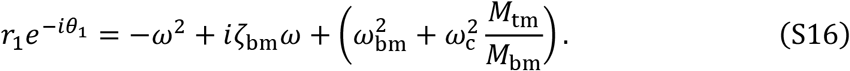

Then (S15) takes form

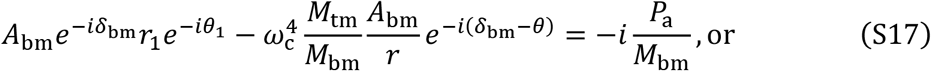

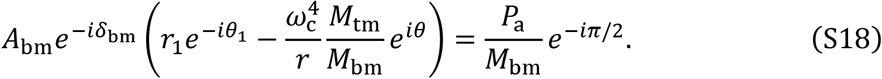

Denote

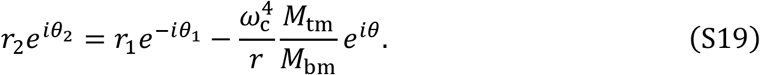

Then (S18) takes form

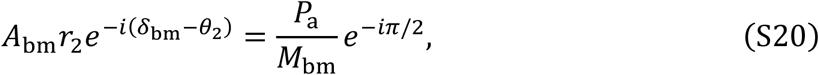

from which it follows that

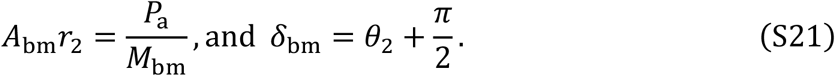

Recall from (S19),

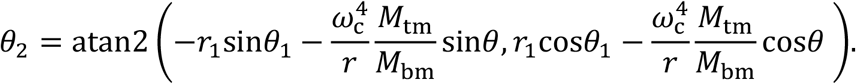

In order to understand behaviour of the phase *θ*_2_, let’s consider the arguments of the *atan*2 function. After substituting all the notations used in the derivation, we have

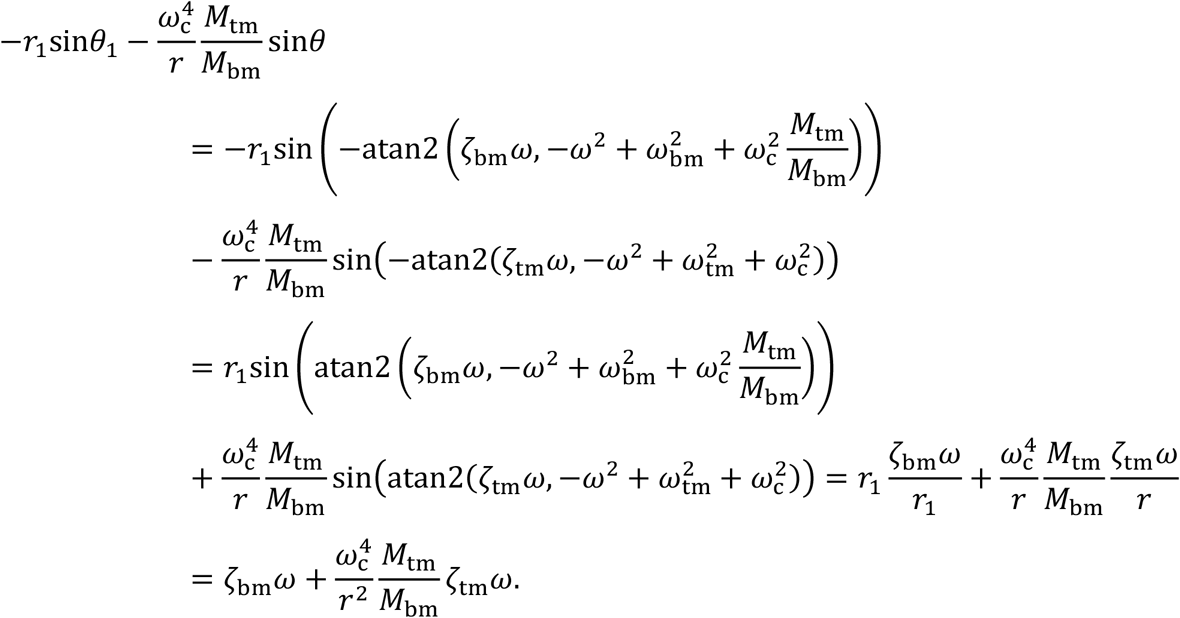

The expression above is positive since *ω* is positive. Note a singularity when *r* = 0 or 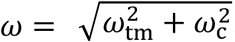, which is where *θ* changes its value from 0 to −*π*.

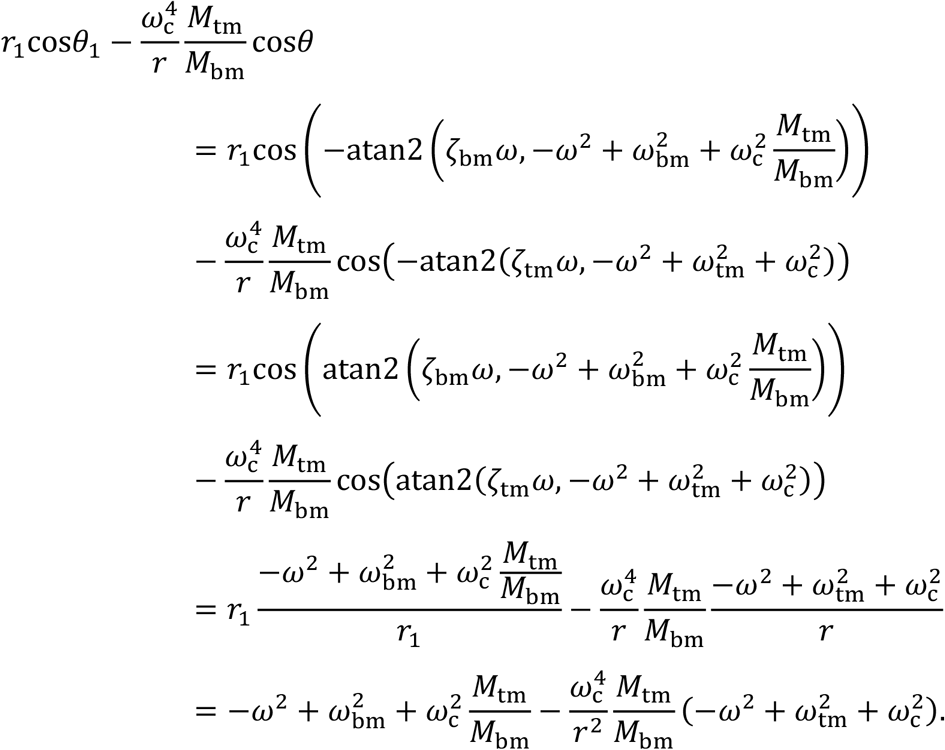

For the purpose of understanding of *θ*_2_, we are interested in the intervals where the expression above is positive or negative intervals. On the positive *ω* semi-axis, there are three points, defining such intervals. Two of them coincide with the system resonances *φ*_1_ and *φ*_2_ (see the derivation of the normal modes in the simplified case, when dampers are neglected below, in the solution to question 2) and 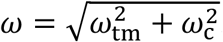. The expression is positive at *ω* = 0. With increasing *ω*, it changes its sign after the first resonance to negative, then becomes positive after 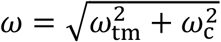 and negative again after the second resonance frequency. If neglecting small terms,

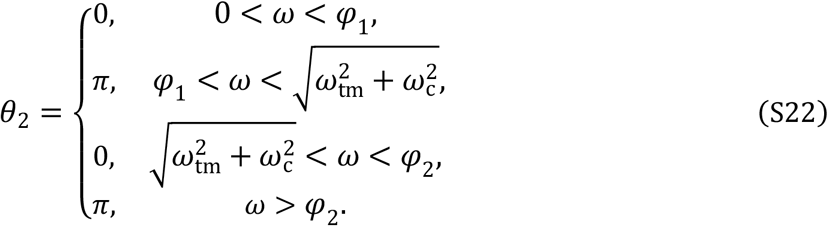

Therefore,

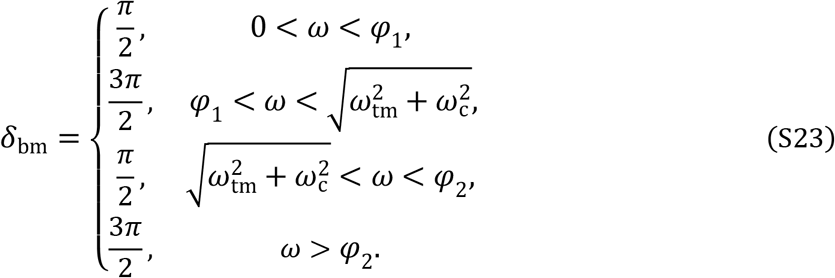

This result is fully replicated in numerical modelling, see Figure 3E, note that *δ*_bm_ has an opposite sign from the BM phase.

Finally, let us consider the difference *Z*_bm_ − *Z*_tm_.

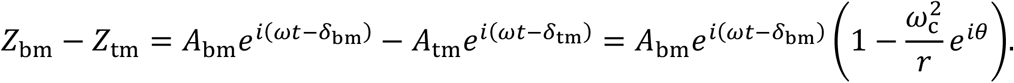

The relative phase of the difference is *Δ*_ph_= *θ*_3_ − *δ*_bm_, where 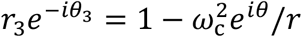. Imaginary part of the latter expression is 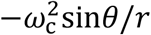. Recall that *θ* is small since it is proportional to 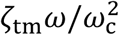. Therefore, *θ*_3_ is close to 0 or *π*, if the real part is positive or negative, correspondingly.

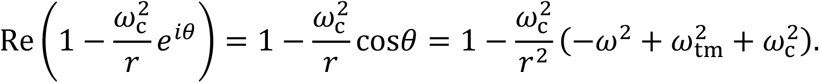

Omitting the details of derivation, there are two points, which divide the positive *ω* semi-axis into three intervals. The first point is close to *ω*_tm_ and the second one is close to the singularity 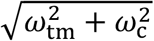. The real part is positive in the first interval between 0 and *ω*_tm_, then changes the sign to negative between *ω*_tm_ and 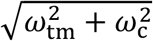, then changes the sign again to positive on 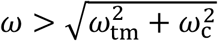. After combining everything together and if neglecting small terms, the phase of the difference takes the following values

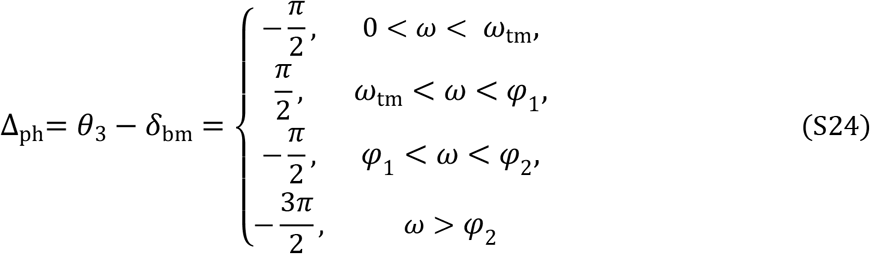

Note that at 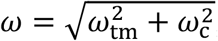, both *δ* and *θ* change their values by −*π*, thus this jump is cancelled out and the value of *Δ*_ph_ remains.

The amplitude of the difference is

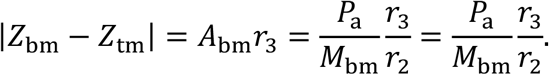

This expression has singularities, which correspond to resonance frequencies (see discussion above regarding *θ*_2_). For the sake of the space, instead of substituting all terms in the above expression, we can focus on the expression for *r*_3_.

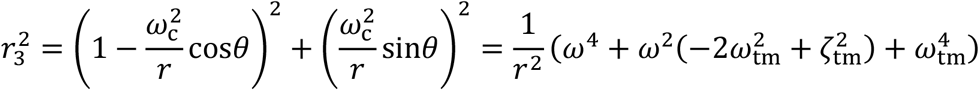

The above shows that since the damping coefficient 0 *< ζ*_tm_ ≪ *ω*_tm_, the minimal value of the amplitude of the difference is when 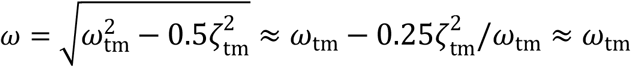, which corrects the previous finding. Note, that at this frequency, 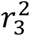 is of the same order of magnitude as 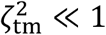.

### Answer to question 1

Minimum of the relative displacement *ΔX* between the TM and BM always occurs at the TM resonance frequency *ω*_tm_ (see also Figure 10 in Nankali et al., 2020). Its frequency position does not depend on the properties of the driven oscillator, i.e. the BM/OoC. The *ΔX* minimum becomes more pronounced with decreasing the TM damping.

To answer question 2, let us consider the original system of equations (4, 5) without dampers and external force to find frequencies of the normal modes:

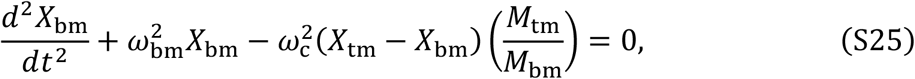

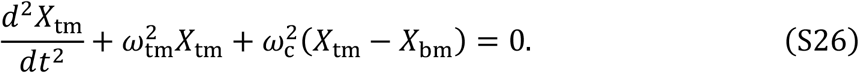

We look for a solution in the form of a cosine function

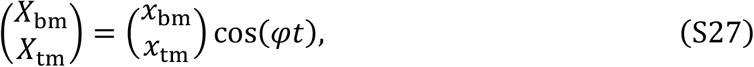

then after substituting (S27) into (S25) and (S26) and using matrix notations we have

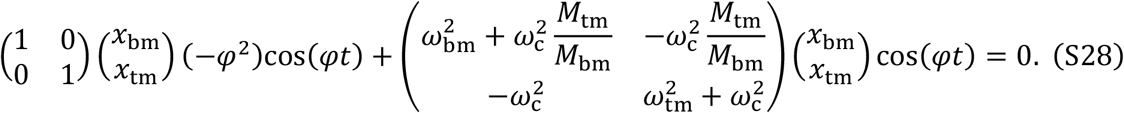

Rearranging yields

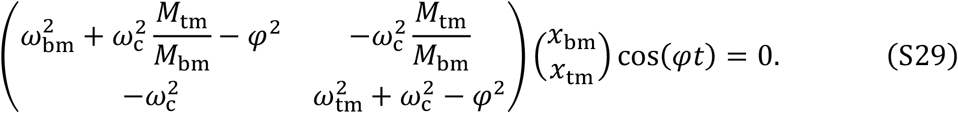

A trivial solution is 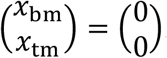. To find non-trivial solutions the determinant of the matrix should be equal to zero, which leads to the following equation with respect to *φ*:

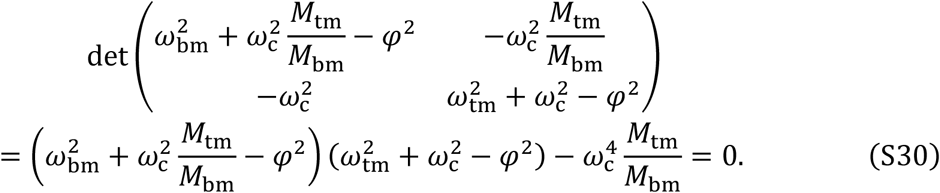

After collecting coefficients of the powers of *φ*, we have

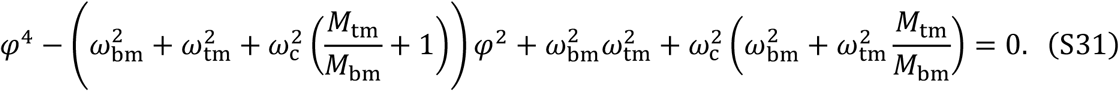

After substituting *Φ* = *φ*^2^, the equation (S31) is reduced to a quadratic equation

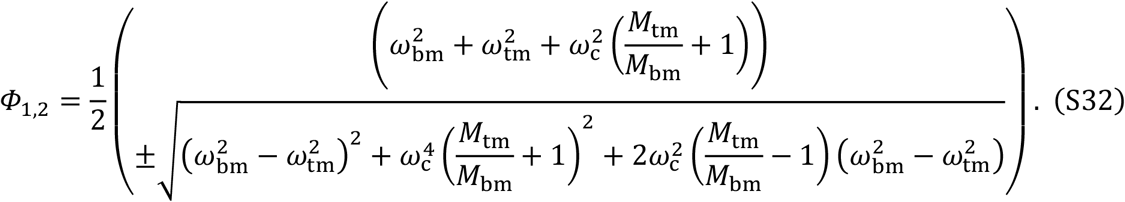

Note that both *Φ*_1_ and *Φ*_2_ are positive and we can find frequencies of the normal modes 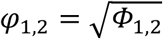. For the chosen model parameters *φ*_1_ = 1.37 and *φ*_2_ = 5.35.

### Answer to question 2

The second normal mode of the system is shifted towards high frequencies due to strong coupling *K*_c_ between the TM and OoC, and its contribution for frequencies below *ω*_bm_ is minimal.

### Answer to question 3

The local minimum of *ΔX* is always observed at *ω*_tm_ (see answer to question 1). Therefore, it is not observed when *K*_tm_ = 0, the TM limbal attachment is absent and *ω*_tm_ = 0. Two normal modes of the system (equation (S32)) exist.

